# Achieving symptom relief in patients with Myalgic encephalomyelitis by targeting the neuro-immune interface and inducing disease tolerance

**DOI:** 10.1101/2020.02.20.958249

**Authors:** Lucie S.T. Rodriguez, Christian Pou, Tadepally Lakshmikanth, Jingdian Zhang, Constantin Habimana Mugabo, Jun Wang, Jaromir Mikes, Axel Olin, Yang Chen, Joanna Rorbach, Jan-Erik Juto, Tie Qiang Li, Per Julin, Petter Brodin

**Affiliations:** Science for Life Laboratory, Department of Women’s and Children’s Health, Karolinska Institutet, SE-17121, SWEDEN; Department of Medical Biochemistry and Biophysics, Karolinska Institutet, SE-17176, SWEDEN; Max Planck Institute Biology of Ageing - Karolinska Institutet Laboratory, Karolinska Institutet, SE-17176, SWEDEN; Department of Clinical Sciences, Intervention and Technology, Karolinska Institutet, SE-17177, SWEDEN; Department of Medical Radiation and Nuclear Medicine, Karolinska University Hospital, SWEDEN; Department of Neurobiology, Care Sciences and Society, Karolinska Institutet, SE-17176, SWEDEN; Neurological Rehabilitation Clinic, Stora Sköndal, Sköndal, Sweden, SE-12864, SWEDEN; Unit of Pediatric Rheumatology, Karolinska University Hospital, SE-17176, SWEDEN

## Abstract

Myalgic encephalomyelitis, ME, previously also known as chronic fatigue syndrome (CFS) is a heterogeneous, debilitating syndrome of unknown etiology responsible for long-lasting disability in millions of patients worldwide. The most well-known symptom of ME is post-exertional malaise, but many patients also experience autonomic dysregulation, cranial nerve dysfunction and signs of immune system activation. Many patients also report a sudden onset of disease following an infection. The brainstem is a suspected focal point in ME pathogenesis and patients with structural impairment to the brainstem often show ME-like symptoms. The brainstem is also where the vagus nerve originates, a critical neuro-immune interface and mediator of the inflammatory reflex which regulate systemic inflammation. Here we report the results of a randomized, placebo-controlled trial using intranasal mechanical stimulation (INMEST) targeting the vagus nuclei, and higher centers in the brain of ME-patients and induce a sustainable, ∼30% reduction in overall symptom scores after eight weeks of treatment. By performing longitudinal, systems-level monitoring of the blood immune system in these patients, we uncover chronic immune activation in ME, as well as immunological correlates of improvement that center around the IL-17 axis, gut-homing immune cells and reduced inflammation. The mechanisms of symptom relief remains to be determined, but transcriptional analyses suggest an upregulation of disease tolerance mechanisms. We wish for these results to bring some hope to patients suffering from ME and inspire researchers to help test our new hypothesis that ME is a condition caused by a failure of inducing disease tolerance upon infection and persistent immune activation.

## Introduction

According to recent estimates ME affects as many as 2.5 million people in the U.S. alone *(1)*. The pathogenesis of ME remains poorly understood and no effective treatment is available for most of these patients. At last, the most recent few years have seen an increased activity in ME research, and the Institute of Medicine has recognized the condition in its recent report as a “serious, chronic, complex system disease” *(2)*, bringing much needed legitimacy to patients facing skepticism and questioning from health care providers. It is now clear that ME has a biological basis, but the disease is highly heterogeneous with variable severity and duration making it challenging to study. One of the most notable characteristic features of ME is the post-exertional malaise leading to significant deterioration of symptoms upon mental or physical activity over an individual threshold that is often quite low *(3)*. An immune system involvement is commonly seen, and many patients present with ME following an acute infection or virus reactivation, and multiple studies have reported elevated markers of inflammation in ME patients *(4)*. Some cytokines correlate with disease severity *(5)* and a recent large study of nearly two hundred ME-patients found differences in T-cell subset composition and function as compared to healthy individuals *(6)*. Our ability to profile human immune systems has improved with the advent of high-dimensional analysis methods such as mass cytometry *(7)*, high-dimensional plasma proteomics *(8)* and single-cell genomics *(9)*. Such methods now allow for all immune cells and many proteins and genes to be captured simultaneously and co-regulated features described *(10)*. Such systems immunology studies in healthy individuals have revealed that human immune systems are incredibly variable among individuals, but very stable within individuals over time *(11)*, and most of this variation is attributed to non-heritable factors *(12)*. Systems-level analyses also provide increased resolution to detect differences in cell, protein and gene combinations associated with disease *(13)*. This becomes apparent when performing longitudinal profiling by systems-level immunomonitoring as perturbations associated with a disease process or response to therapy become apparent, and often such patterns would not be visible from reductionist analyses of individual immune system components alone *(14–16)*.

Another clinical feature of ME is autonomic dysregulation, i.e. dysautonomia, with impairments to blood pressure regulation, temperature regulation, intestinal function and a range of other central body functions. Given the concomitant immune and autonomic perturbations in ME it is conceivable that a common culprit could involve the neuro-immune interface. The most notable such interface is the inflammatory reflex, a well-known anti-inflammatory reflex mediated by the vagus nerve *(17)*. Direct infection of vagus nerve fibers has been suggested as a possible pathogenic mechanism in ME *(18)* and some patients with cervical compression involving the vagus nucleus present with ME-like symptoms, sometimes reversible upon decompressive surgery *(19)*. Moreover, afferent fibers in the vagus nerve can sense microbial stimuli and dysbiosis in the gut and elsewhere that fuel the chronic inflammation in ME *(20)*.

Several treatment modalities target the vagus nerve; implanted electrical nerve stimulators in patients with severe epilepsy, as well as non-invasive stimulators using mechanical stimulation of mechano-sensitive nerve endings in the ear *(21)*, or on the neck *(22)*, can stimulate the vagus nerve nuclei in the brainstem and trigger the inflammatory reflex. Alternative treatment modalities, Intranasal Mechanical Stimulation (INMEST) and the related Kinetic Oscillatory Treatment (KOS), use mechano-stimulation to trigger nerve endings in the nasal cavity, that propagate signals to the vagal nerve nuclei in the brain stem *(23, 24)*. Curiously, the use of these treatments in patients with acute migraine and chronic inflammatory disorders have indicated a dampening of systemic inflammation *(25, 26)*. Functional MRI in healthy volunteers treated with INMEST show increased activity in the vagus nerve nuclei *(24)*, but the effects on heart-rate variability, a complex measurement of autonomic regulation and the balance between sympathetic and parasympathetic tone, differs between these vagus stimulating treatment modalities *(23)*.

Here we reasoned that using INMEST in ME could induce symptom relief similar to patients with other inflammatory disorders *(26)*. Also, by performing longitudinal systems-level immunomonitoring in treated patients, our hope was to better understand the pathways perturbed in ME by monitoring changes associated with treatment responses and possible symptom relief. We performed a placebo-controlled, double-blinded, randomized trial of INMEST given biweekly for 8 weeks. The clinical improvement was significantly better with active INMEST than with placebo and represented ∼30% reduction in symptom intensity after 8 weeks. We also found measurable changes in immune measurements that suggest perturbations involving IL-17-mediated inflammation, immune cell energy metabolism and a failure to induce disease tolerance as elements of ME pathogenesis. These results suggest novel lines of mechanistic investigation into disease and that INMEST is a simple and safe method for symptom relief through a re-calibration of the neuro-immune setpoint.

## Results

### ME Symptom relief after INMEST treatment

We enrolled 31 patients with moderate to severe ME (17 in 2018, and 14 in 2019). All of these fulfilled the Canadian consensus criteria. These subjects were randomized to one of two arms within a double-blinded, randomized control trial to receive 20 minutes of INMEST twice a week for 1 month, or placebo treatment, which is indistinguishable from active INMEST to the patient and the treating physician. This point is important to avoid bias from changes to the patient-doctor relationship. Following the first month of placebo or active treatment, all subjects were exposed to active INMEST treatment 20 minutes bi-weekly (**Fig. 1A**). The INMEST device consists of a thin plastic probe placed in the nose that vibrates at a set frequency to mimic turbulent airflow within the nasal cavity and induces a nerve reflex transmitted to the vagus nerve nuclei in the brainstem and other higher centers (24) (**Fig. 1B**). It is important to note that the effect of INMEST on the vagus nerve is different from that of classic vagus nerve stimulation. INMEST shifts the balance of sympathetic and parasympathetic tone resulting in a reduction of heart rate variability (23).

**Fig. 1.**
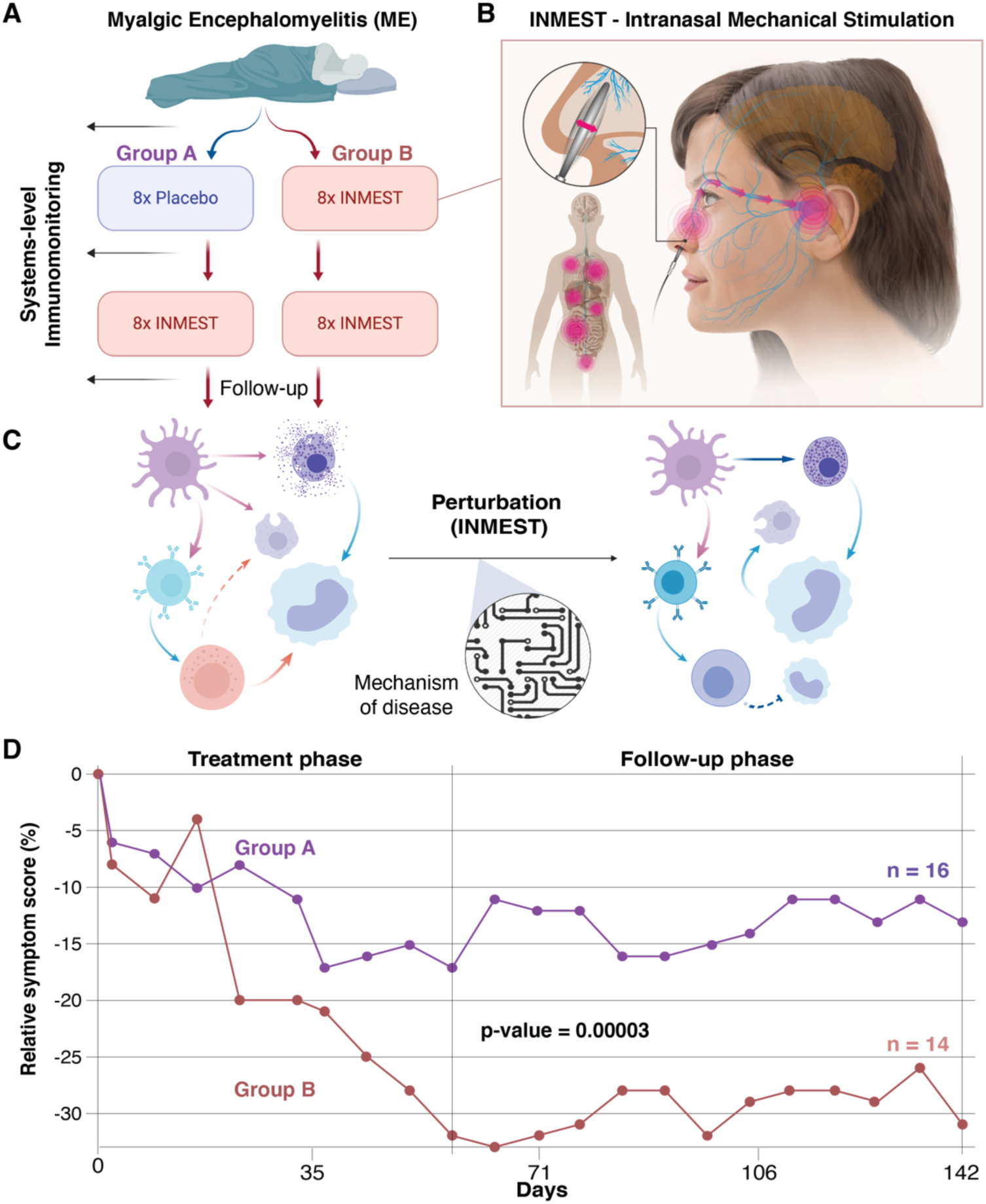
Study design, treatment and outcome. **(A)** Study design and sample collection for systems-level immunomonitoring. **(B)** Intranasal Mechanical Stimulation, INMEST. **(C)** Treatment (INMEST) is used as a perturbation to probe mechanisms of disease in ME-patients. **(D)** Change in symptom score in patients randomized to Group A vs. Group B. Y-axis indicate average ME symptom scores relative to baseline.

This study is not powered to test the possible beneficial effects of the INMEST treatment on patient outcome. Instead, the purpose of the INMEST treatment here is to induce a perturbation to the immune system, that in combination with systems-level monitoring will allow us to decipher the mechanisms of immune system perturbation in ME-patients (**Fig. 1C**). This method of system-perturbation and longitudinal monitoring is common in systems biology and an powerful tool to better understand biological pathways and systems (27).

The primary outcome measure in this study was fatigue as defined by the fatigue severity scale (FSS) (28), but no significant effect of INMEST treatment was seen with respect to this outcome variable. We also evaluated overall symptom scores using a questionnaire previously shown to distinguish subgroups of ME-patients (29). During the first month of treatment, the placebo showed a modest response of ∼10% reduction in symptom score, while the actively treated patients initially improved, then displayed a rebound effect, followed by a more significant improvement and a total ∼20% overall reduction in symptom scores after the first four weeks (8 active treatments) (**Fig. 1D**). It is important to note that many patients struggled with biweekly visits to the clinic and some patients suffered episodes of deterioration as a consequence of post-exertional malaise. During the second phase of active INMEST treatment (for all patients), symptom scores decreased further and in patients receiving active treatment only ∼30% decrease in symptom scores were achieved at the end of the 8-week trial (16 active treatments) (**Fig. 1D**). Since treatments varied slightly over different days, but result was the same if plotted against number of treatments rather than days of study (**Fig. 1D**). The effect was significantly stronger (p=0.00003) in the patients receiving active treatment only vs. patients receiving placebo followed by active treatment (**Fig. 1D**), and the response persisted after cessation of treatment in both groups and even improved further in the most actively treated group, suggesting that further improvements could be possible with a prolonged treatment protocol (**Fig. 1D**).

### Immune system measurements correlate with symptom relief

To understand the underlying immune system perturbation in ME, and investigate any potential changes correlating with treatment response, we collected blood samples at baseline before treatment, after phase 1 (4 weeks), and after phase 2 (8 weeks) (**Fig. 2A**). To minimize technical variation, samples were prepared directly at blood draw and cells frozen in whole blood stabilizer solution (30), plasma centrifuged and frozen, and whole blood bulk mRNA stabilized and frozen (31) (**Fig. 2A**). We performed immune cell compositional and phenotypic analysis using mass cytometry, plasma protein analysis by Olink assays and whole blood mRNA-sequencing to gather different perspectives of the global immune state in ME-patients before, during and after treatment time. The gene most significantly induced by active INMEST treatment was the E2 ubiquitin-conjugating enzyme UBE2H (**Fig. 2B**), a gene broadly expressed by blood immune cells (https://www.proteinatlas.org/ENSG00000186591-UBE2H/blood) that stimulates ubiquitin mediated degradation of dysfunctional proteins.

**Fig. 2.**
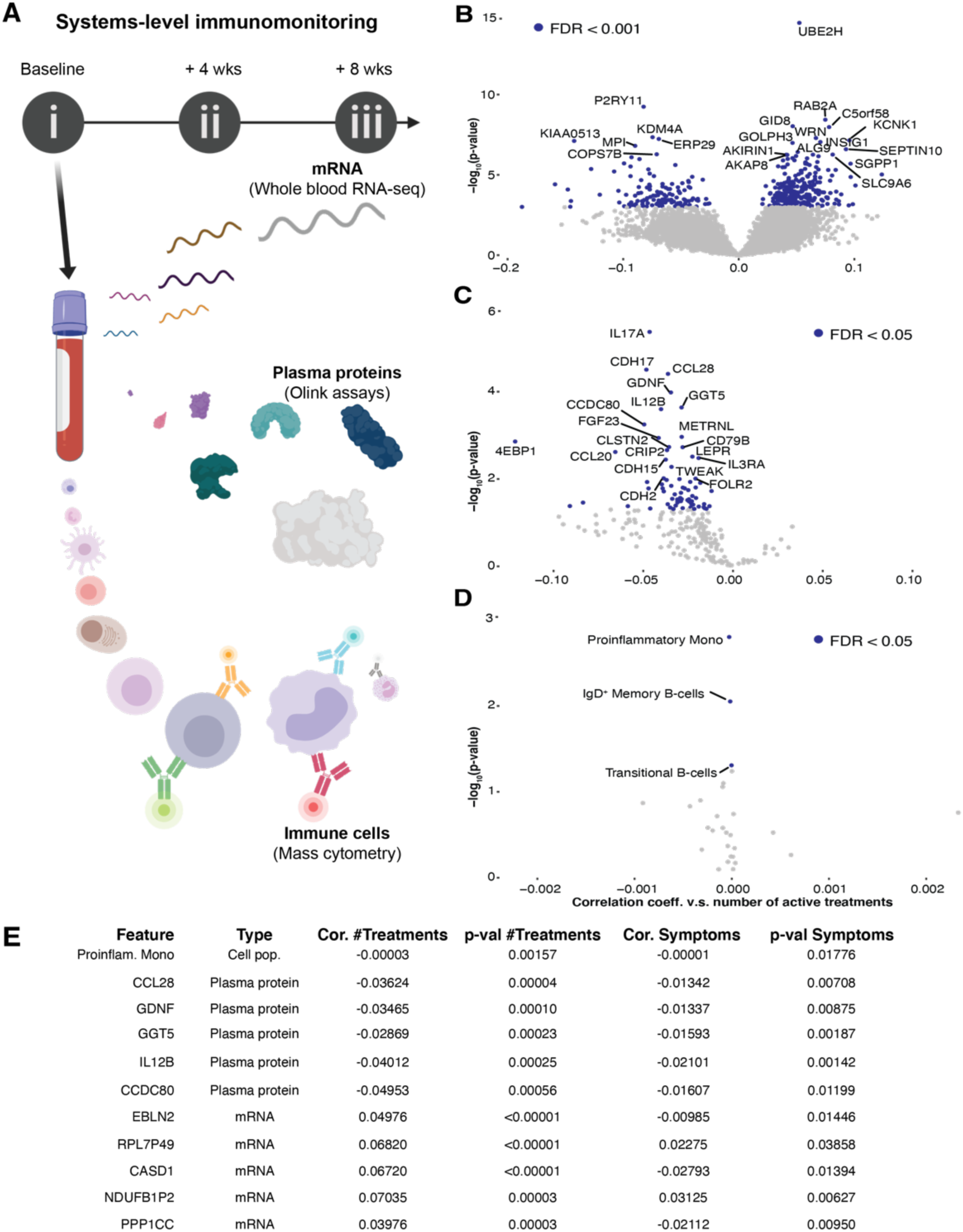
Systems-level profiling reveal immune perturbations. **(A)** Blood samples were collected and immediately processed for whole blood mRNA-seq (PaxGene tubes), Mass cytometry (Stabilized whole blood cells) and plasma proteins (immediate centrifugation) by Olink assays. **(B-D)** Volcano plots of differentially regulated features v.s number of active treatments. **(E)** Mixed effects modeling shows the most perturbed plasma proteins and immune cells that change in response to therapy and in relation to symptom scores.

Plasma cytokines IL-17A, IL-12B and CCL28 were among the top plasma proteins decreasing after INMEST treatment (**Fig. 2C**) and these have all been previously reported to be involved in ME (32). Circulating immune cells also changed, especially the proinflammatory monocytes that decrease over time during INMEST treatment (**Fig. 2D**).

Next we used mixed-effects modeling to test whether individual immune system measurements were associated with symptom relief as measured by symptom score questionnaires. We also used age, sex, treatment group (pre-/post-INMEST) and number of treatments as fixed effects and found that frequencies of proinflammatory monocyte in the blood, plasma IL-12B and CCL28 and other pro-inflammatory mediators decreased as symptom scores decreased after INMEST treatment (**Fig. 2E**). These results further underscore the inflammatory nature of ME (1) with variable levels of cytokine elevation in plasma that correlate with symptom severity. Our results also show that the biomarker correlates of response in ME patients can be measured in the blood.

### Multi-Omics Factor Analysis reveals co-regulated biomarkers in ME

Immune cells and proteins do not function in isolation but coordinate their actions into a complex network of communicating features. Multi-Omics Factor Analysis (MOFA) is a method for integrating different types of data and allow such coordinated features to be identified and associated with a condition of interest in the form of latent variables that discriminate groups of samples (32). MOFA was applied in our cohort to all plasma proteins, immune cell frequency and mRNA-measurements across all blood samples (n=85). A range of variable MOFA models were run with different constraints until a model with 10 latent factors (LFs was found to explain the variance among the samples (**Fig. 3A-B**). We inspected samples across these LFs to identify any factor that could segregate samples collected before and after INMEST treatment. LF7 showed such a gradient from pre-to post-INMEST samples and we focused on this latent factor (**Fig. 3C**). In LF7 65% of genes and 50% of plasma proteins overlapped with the significant features in the mixed-effect model above, emphasizing that features associated with INMEST response can be measured in blood, but the MOFA result also broadens the scope of immunological features influenced by INMEST and also suggests a shift from CD56^dim^ to more CD56^bright^ type of NK-cells, known to be immunoregulatory cytokine-secreting cells (**Fig. 3D**) (33).

**Fig. 3.**
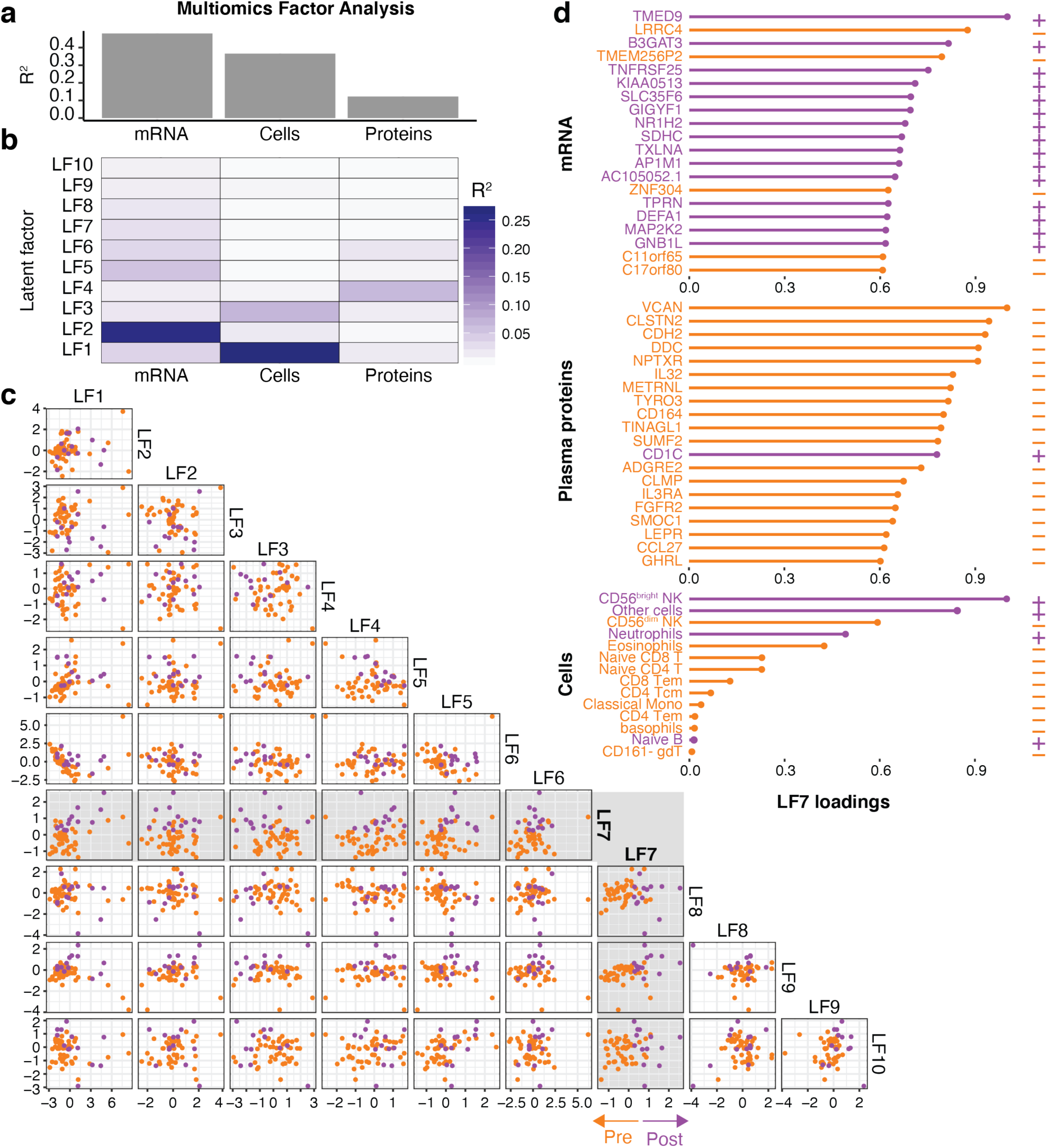
Co-regulated immune features revealed by multi-omics factor analyses. (**A**) Variance explained by different views (Immune cells, proteins and mRNA), (**B**) Variance explained by each of the calculated latent factors, (**C**) Scatterplot showing individual samples and their latent factor distributions, (**D**) Top features contributing to latent factor 7 associated with INMEST treatment response.

### Cell-cell relationships are perturbed in ME

Given the coordinated changes induced by INMEST-treatment across different immune system components, we decided to take a closer look at cell-cell dependencies and how these would be rewired by treatment. This is important because all immune responses occur as concerted efforts by multiple cell populations, and perturbations to such relationships can be an element of disease (34). To this end, we compared cell-cell correlation matrices before and after INMEST treatment and we found specific cell-cell relationships changing after INMEST treatment (**Fig. 4**). For example, positive associations were found between naïve and central memory T-cells, that became more pronounced after INMEST, likely reflecting an improved cell-cell coordination within the system as a consequence of dampened inflammation described above (**Fig. 4B**).

**Fig. 4.**
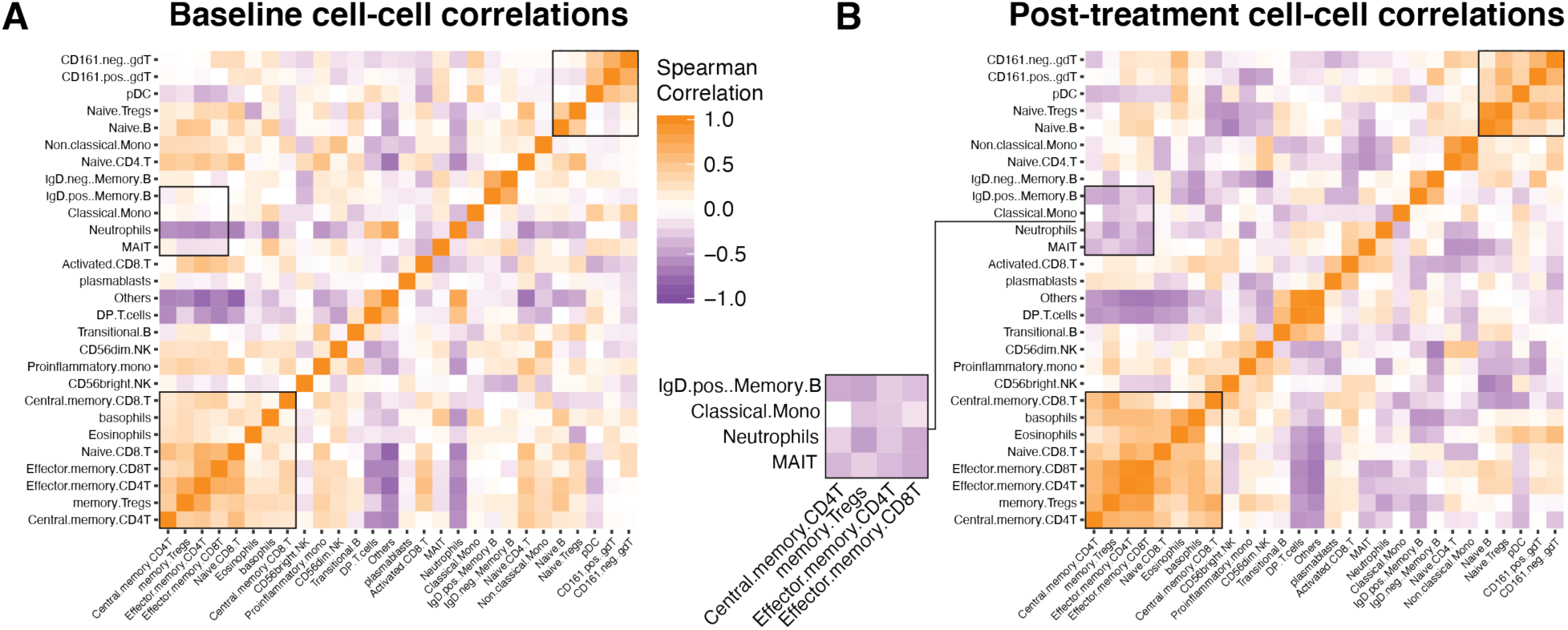
Immune cell function in ME patients treated with INMEST. **(A)** Spearman correlation matrix showing co-regulated cell population frequencies at baseline, and **(B)** After 16 active INMEST treatments, indicating re-establishment of specific cell-cell relationships, such as the highlighted module involving MAIT-cells and marginal zone B-cells (lgD+ mem. B) negatively correlated with memory Tregs.

Moreover, a specific module of cells involved Mucosal-associated invariant T-cells (MAIT cells) and marginal zone (IgD^+^ memory) B-cells, both known to be important cells interacting with microbes in the gut (18). MAIT-cells have been implicated in ME previously (19)(6). In these cell-cell correlation analyses, these effector cells correlated inversely with antigen-experienced (memory) Regulatory T-cells, a type of cell known for its critical importance in maintaining homeostasis at barrier surfaces such as the gut (35). This relationship was not visible at baseline (**Fig. 4A**), but appeared after INMEST treatment, indicating a normalization of a critical immune cell module and regulatory mechanism in parallel with symptom relief (**Fig. 4B**). We think this finding represents another clue into ME-pathogenesis and the mechanism of action of INMEST, since the gut microbes and dysbiosis is likely in ME (20), and the vagus nerve conveying such afferent signals from the gut to the brainstem and inducing immune regulation via the inflammatory reflex (*27, 28*). The reduction in plasma IL-17A and CCL28 levels after INMEST and the restored MAIT/Regulatory T-cell relationship indicate a modulation of this axis by INMEST treatment and mitigation of an inflammatory state originating at the immune microbe interface in the intestine. Also, the common Epstein-Barr virus, EBV, often associated with ME have been reported to induce IL-17 and reactivation of this virus is another possible chronic immune stimulation in ME-patients (36).

### Transcriptional programs altered in ME patients treated by INMEST

To further investigate the possible mechanisms of action of INMEST in these ME-patients, we investigated the whole blood transcriptome more in-depth. INMEST treatment upregulated 142 genes (p-value < 0.05) and downregulated 12,384 genes (p-value < 0.05), a gene set enrichment analysis (30) was performed on these differentially regulated genes in order to understand the molecular pathways affected (**Fig. 5A**). The most enriched Gene Ontologies (GO) are shown in Figure 4A and involve three broad biological processes; I) nerve cell signal transmission, II) immune response pathways and III) cellular metabolism (**Fig. 5A**), all previously linked to the pathogenesis of ME (1).

**Fig. 5.**
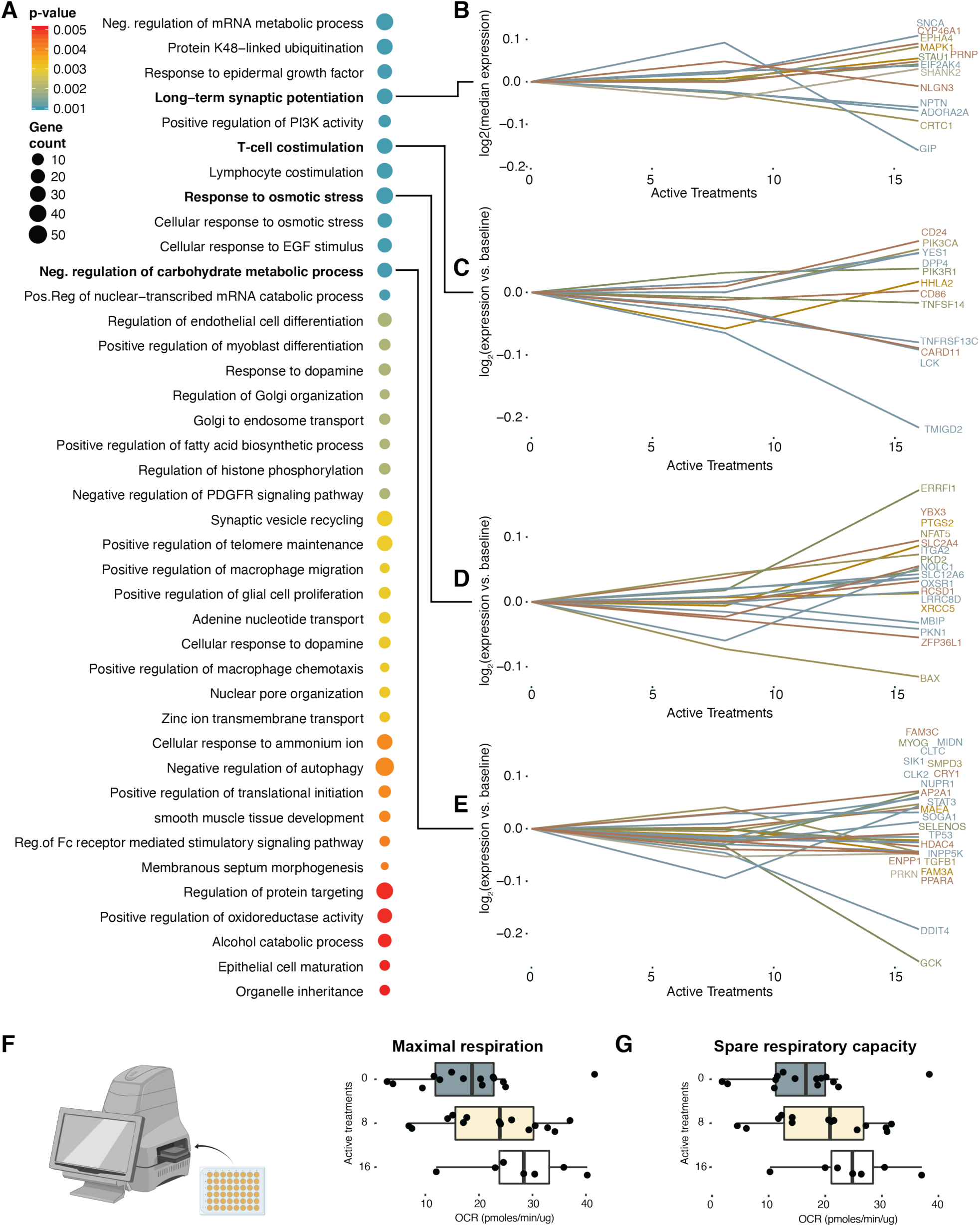
Immune cell metabolism is impaired in ME/CFS but normalized by INMEST. **(A)** Gene Set Enrichment Analysis, GSEA for genes differentially regulated in response to INMEST treatment. **(B-E)** Four highlighted GO terms and the top genes with the largest absolute change in response to INMEST treatment. **(F)** Maximal respiration in cultured PBMC from ME/CFS patients at baseline, and upon 8 and 16 active INMEST treatment rounds respectively (n = 3). **(G)** Spare respiratory capacity, SRC in cultured PBMC from ME/CFS patients at baseline, and upon 8 and 16 active INMEST treatment rounds respectively (n = 3).

A set of genes involved in *Long-term synaptic potentiation* increased with INMEST treatment, particularly after >8 treatment sessions (**Fig. 5B**). Examples include SYNC, a gene encoding the protein alpha-synuclein present in neuronal presynaptic terminals and regulating neuronal signal transmission (37). The T-cell co-stimulation gene set involves molecules such as CD86 on antigen-presenting cells, which are necessary for co-stimulating T-cells activated by antigens. Most of these genes decreased after INMEST in line with an overall dampening of immune cell activation in ME-patients treated with INMEST (**Fig. 5C**).

In a recent study by Davis and colleagues, blood cells from ME patients were reported to display a unique impedance pattern in response to hyperosmotic stress suggesting that a nano-electronic impedance sensor could be used in a diagnostic test for ME (38). We find that the GO: *Response to osmotic stress* was associated with INMEST treatment, and genes involved include the Bcl2 family member BAX, upon treatment (**Fig. 5D**).

Finally, multiple gene sets regulated by INMEST involved cellular energy metabolism, which is curious given that fatigue is the cardinal symptom of ME and linked to alterations in cellular metabolism (39). *The GO: Negative regulation of cellular carbohydrate metabolism* is affected by INMEST with genes like GCK encoding the glucose sensor Glucokinase, which shifts cellular metabolism based on the availability of glucose (40) and was found in this cohort to be repressed in ME-patients after INMEST treatment (**Fig. 5E**).

To investigate changes in energy metabolism in ME patients, we took advantage of the Agilent Seahorse analysis energy metabolism using cultured PBMCs collected from ME-patients at baseline, after 8 and 16 treatment rounds, respectively (**Fig. 5F**). This assay measures multiple aspects of cellular metabolism and ATP usage and oxidative phosphorylation. As predicted by mRNA-sequencing results above, we find improvements in energy metabolism in PBMCs from ME-patients treated with INMEST, both in Maximal respiration (**Fig. 5F**), and spare respiratory capacity (**Fig. 5G**), suggesting that INMEST impacts multiple pathways previously reported to be perturbed in ME.

### ME - a failure of inducing disease tolerance upon chronic immune activation

Given the evidence of chronic immune activation in ME and the multifaceted symptoms in ME we considered possible unifying mechanisms that could explain the variable presentation of this mysterious illness. Disease tolerance is an overall term describing a range of stress-response pathways that limit tissue damage caused by invading pathogens or indirectly by host immune responses (41). We hypothesized that ME could be due to a failure of upregulating such disease tolerance mechanisms in response to infection. To this end, we analyzed genes involved in disease tolerance induction based on a transcriptional regulatory network database TTRUST v (42). Most effector genes increased their mRNA expression after active INMEST treatments in ME-patients (**Fig. 6A-C**), supporting the hypothesis that induction of disease tolerance could mitigate ME symptoms. We then focused on disease tolerance pathways most strongly upregulated with treatment. HIF-regulated effector genes like VEGFA, CXCR4, AR and SERPINE1 are induced by tissue hypoxia during infections, a response important for limiting tissue damage potentiated by INMEST treatment (**Fig. 6A-C**). Ramasubramanian et al has reported higher levels of reactive oxygen species (ROS) in red blood cells of ME-patients compared to healthy controls (43), and up-regulation of the Oxidative stress response pathway upon INMEST (SIRT1), corroborates this observation. These upregulated pathways and the induction of genes in the insulin receptor signaling pathway, Heat shock proteins, and cell cycle regulators all support the idea that INMEST treatment upregulates disease tolerance pathways normally induced during infections, and possibly improperly suppressed in ME-patients (**Fig. 6A-C**). In conclusion our findings show evidence of chronic immune activation in patients with ME, particularly involving the IL-17 pathway, MAIT-cells and other intestinal lymphocytes interacting with microbes. The INMEST treatment affecting via the vagus nerve induces a significant symptom relief that correlates with a normalization of immune cell regulatory networks, cellular metabolism and upregulation of disease tolerance programs.

**Fig. 6.**
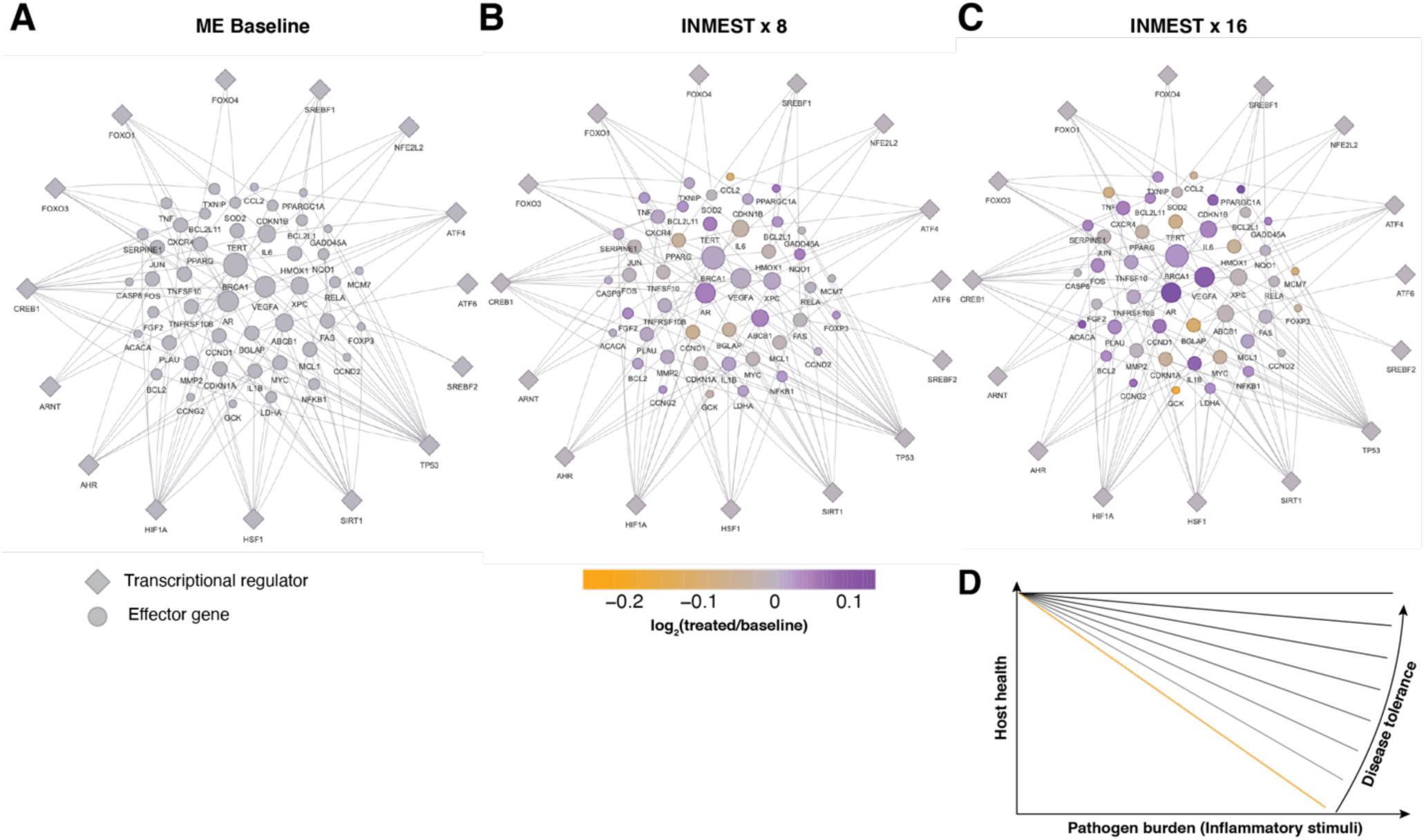
INMEST treatment induces disease tolerance programs in ME-patients. Transcriptional regulators (diamonds) and their target genes involved in disease tolerance are shown in ME-patients **(A)** at baseline and **(B)** after 8- and **(C)** 16-rounds of INMEST-treatment. **(D)** A proposed model of ME as a disease caused by failure to upregulate disease tolerance mechanisms in response to chronic inflammatory stimulus leading to broad impairments and reductions in overall health. Size of the nodes refers to number of regulatory interactions (larger = more regulatory interactions)

## Discussion

Our findings paint a picture of ME as a truly multifaceted disorder involving inflammation with a possible origin at the immune-microbe interface in the gut. Chronic immune stimulation is likely given the symptomatology, but also elevated cytokine levels are found that correlate with symptom severity (5) and immune cell deficiencies (6). The deficiencies in cellular metabolism are not typically seen in other inflammatory diseases, although transient metabolic adaptations are common during immune responses (44).

It is important to note that ME is a heterogeneous disease and the likelihood of finding one pathogenic mechanism shared by all patients is low. We believe that one unifying concept could be the failure in upregulating disease tolerance mechanisms in the event of an infection or virus reactivation and this is what we propose herein. Such disease tolerance mechanisms are important in limiting the reduction in host fitness as a consequence of the infection directly or as a consequence of the elicited immune defenses (41). Our data of upregulated disease tolerance pathways upon INMEST treatment support the hypothesis that ME is a result of a failure to upregulate disease tolerance mechanism when faced with infection and thereby leading to a deterioration of physiological functions (**Fig. 6D**), although the precise mechanisms of this remains to be unraveled. The reasons why ME patients differ so much in their symptomatology could be explained by differences in the underlying infectious disease or immune activation, and the relevant disease tolerance pathway failing to be induced. The reasons for why ME patients would fail to upregulate disease tolerance pathways remains to be determined, although one interesting lead comes from an observation made of mutations in the enzyme IDO2 seen in 20/20 patients with severe ME (https://www.omf.ngo/2018/10/19/healthrising-themetabolic-trap-shines-during-the-symposium-on-the-molecular-basis-of-me-cfs-at-stanford/) and has led to the formulation of the metabolic trap hypothesis of ME (45). However, the IDO-enzymes are also involved in disease tolerance, specifically upon exposure to Endotoxin, a component of Gram-negative bacteria in the gut (46, 47).

The symptom relief induced by INMEST targeting the vagus nerve is significant and distinct from placebo, but must still be confirmed in larger trials with sufficient statistical power. We believe that this should be done using a self-treatment system available for use at home since the repeated visits to the clinic are so demanding for patients with ME. The main purpose of the current study was instead to use the INMEST treatment as a perturbation to the immune system and autonomic inflammatory reflex, as a means of uncovering the pathogenesis of the disorder. To this end the current study was successful and the biomolecular correlates found corroborate several previously suggested aspects of ME pathogenesis.

The mechanism of action of the INMEST-treatment in ME is not known, although some things are clear. We know that the vagus nerve nucleus in the brainstem is activated by INMEST, but also higher level centers such as the limbic system are activated (24). The effect of INMEST on heart rate variability differs from that of traditional vagus nerve stimulating methods (23). One possibility is that INMEST influences incoming (afferent) signals from the gut, conveying signals of dysbiosis or chronic immune activation and inflammation. This hypothesis is in line with previous proposals of ME as a disease caused by microbial dysbiosis in the gut (48). One possible mechanism of symptom relief upon INMEST-treatment could be through limiting such signals of enteric dysbiosis via the afferent vagus nerve.

A problem with the treatment trial presented here is that ME-patients were forced to visit the clinic twice weekly for 8 weeks to be treated and blood sampled as well as respond to questionnaires. All of these activities were very demanding and caused deterioration in some patients that obscure some of the possible benefits of the therapy. We found no significant improvement of the primary outcome measurement, fatigue in this study. One possible explanation for this is that the recurrent clinical visits and the consequent worsening of disease. Another possible explanation is related to the Fatigue Severity Scale, FSS used because. This questionnaire has been shown to best distinguish patients with ME from healthy individuals, while its ability to capture quantitative changes within patients over time is in question, and ceiling effects have been reported (49). In the future a more sustainable way of using INMEST would be in the form of a self-treatment device used by subjects and care takers at home. Obviously larger follow-up studies are required to further assess the clinical value of this approach more generally in ME-patients. It is likely important to improve the diagnostic accuracy of ME and ensure more homogenous group of patients included in order to sharpen our ability to understand the underlying pathology in ME, subdivide patient groups and test treatment strategies in these different subgroups accordingly. The recent advances in diagnostic tests are promising in this regard (38). We hope that this work will inspire others to further investigate the immune/microbe interface and the failure to induce disease tolerance in ME and hopefully this increased attention will lead to some further relief to the millions missing due to this devastating condition.

## Materials and Methods

### Study Design

The sample size was based on the patients who had volunteered to take part in the clinical trial up to the point we started analysis, therefore both of the first two cohorts were included in the study. All samples were included in the trial unless the patients opted to drop-out of the clinical trial before the end of the study. The Swedish ME cohort that underwent the INMEST treatment comprises of a gender-mixed group, 23 females and 8 males. Female patients had a median age of 42 (range 19-62) and male patients had a median age of 35 (range 26-66). All were evaluated for enlistment based on the Canadian consensus criteria *(49)*. The INMEST treatment for the clinical trial (NCT03502044) was undergone at the Neurological Rehabilitation Clinic in Stora Sköndal with Dr. Per Julin as the lead investigator. The research group has used INMEST and KOS interchangeably until 2018, reason for the latter term being used on clinicaltrials.gov.

### Experimental Design

The randomized, placebo-controlled, and double-blinded trial of intranasal mechanical stimulation (INMEST) included two weekly doses over the span of 8 weeks. The doses included 10 min of placebo or active INMEST treatment in each nasal cavity (20 min total). For each cohort, the INMEST device randomly selected either placebo or active treatment during the first 4 weeks followed by only active treatments the following 4 weeks. During the first 4 weeks both the patient and doctor were blind as to whether placebo or active treatment was given. In the end, the active-active group included 5 males and 11 females while the placebo-active group included 3 males and 12 females.

One of the key factors that was recorded during this study is the ME symptom rating scale, it is a rating scale of the severity of symptoms from the ICC criteria on a 5 degree scale from 0-4 (none, light, moderate, severe, very severe) in order to evaluate degree of disease burden in accordance to the Canadian Consensus criteria. Other than symptom severity, the following measures were recorded for the clinical trial: Fatigue severity scale, SF-36 Physical functioning subscale (PF-10), Hospital anxiety depression scale, EQ5D, as well as VAS. Blood samples were collected at 3 timepoints for each patient (baseline and subsequentially either Pre-/Post-treatment depending on placebo or active group meaning 8 or 16 active treatment).

### Immune cell phenotyping by Mass Cytometry

Cryopreserved and stabilized whole blood (blood mixed with ‘Stabilizer’ component of Whole blood processing kit; Cytodelics AB, Sweden) collected from ME patients sampled thrice (Baseline and T1 [4 weeks] and T2 [8 weeks]) during the study period were thawed, and cells were fixed and RBCs lysed using lysis and wash buffers (Whole blood processing kit; Cytodelics AB, Sweden) as per the manufacturer’s recommendations. This was performed a few days prior to barcoding and staining of cells. Post fix/lysis of cells, ∼1×10^6^ cells/sample were plated onto a 96 well ‘U’ bottom plate using standard cryoprotective solution (10% DMSO and 90% FBS) and cryopreserved at −80°C. On the day of barcoding and staining of cells, cells were thawed at 37°C using RPMI medium supplemented with 10% fetal bovine serum (FBS), 1% penicillin-streptomycin and benzonase (Sigma-Aldrich, Sweden). Briefly, cells were barcoded using automated liquid handling robotic system (Agilent Technologies, Santa Clara, CA, USA)*(38)* using the Cell-ID 20-plex Barcoding kit (Fluidigm Inc.) as per the manufacturer’s recommendations. Following cell pooling batch-wise (with samples from placebo and treatment groups equally represented in each batch), cells were washed, FcR blocked using blocking buffer (in-house developed recipe) for 10 min at room temperature, following which cells were incubated for another 30 min at 4°C after addition of a cocktail of metal conjugated antibodies targeting the surface antigens. Following two washes with CyFACS buffer, cells were fixed overnight using 4% formaldehyde made in PBS (VWR, Sweden). The broad extended panel of antibodies used for staining are listed in Supplementary Table 1. For acquisition by CyTOF (within 2 days after staining), cells were stained with DNA intercalator (0.125 μM Iridium-191/193 or MaxPar® Intercalator-Ir, Fluidigm) in 4% formaldehyde made in PBS for 20 min at room temperature. After multiple washes with CyFACS, PBS and milliQ water, cells were filtered through a 35µm nylon mesh and diluted to 750,000 cells/ml. Cells were acquired at a rate of 300-500 cells/s using a super sampler (Victorian Airship, USA) connected to a CyTOF2 (Fluidigm) mass cytometer, CyTOF software version 6.0.626 with noise reduction, a lower convolution threshold of 200, event length limits of 10-150 pushes and a sigma value of 3 and flow rate of 0.045 ml/min.

### Antibodies and reagents

Purified antibodies for mass cytometry were obtained in carrier/protein-free buffer and then coupled to lanthanide metals using the MaxPar antibody conjugation kit (Fluidigm Inc.) as per the manufacturer’s recommendations. Following the protein concentration determination by measurement of absorbance at 280 nm on a nanodrop, the metal-labeled antibodies were diluted in Candor PBS Antibody Stabilization solution (Candor Bioscience, Germany) for long-term storage at 4°C. Antibodies used are listed in Supplementary Table 1.

### ProSeek data collection

Plasma protein data was generated using the proximity extension assay (ProSeek, Olink AB, Uppsala). Three panels (Inflammation, Metabolism and Neuro-Exploratory) were used to detect a range of biomarkers. Each panel kit provides a microtiter plate for measuring 92 protein biomarkers. Each well contains 96 pairs of DNA-labeled antibody probes. Samples were incubated in the presence of proximity antibody pairs tagged with DNA reporter molecules. When the antibody pair bounds to their corresponding antigens, the corresponding DNA tails form an amplicon by proximity extension, which can be quantified by high-throughput real-time PCR. The data was generated in a single batch (90 samples).

### Whole blood mRNA-sequencing

Two milliliters of whole blood cryopreserved in PAXgene Blood RNA Tubes (cat.nr. 762165, Qiagen) were subjected to RNA isolation with QIAcube using PAXgene Blood RNA Kit (cat.nr. 762164, Qiagen). After elution, RNA quantity and quality (RIN value) were inspected with Qubit RNA HS Assay Kit (ThermoFisher, cat.nr. Q32855) and Agilent Technologies, cat.nr. 5067-1513), respectively.

Initial steps of RNA isolation consist of cell washing with ultrapure water followed by centrifugation to remove red blood cells. However, we could detect the presence of erythrocytes contents after washings (red pellet and high presence of mRNA from erythrocytes confirmed by qPCR). This interference of transcripts from red blood cells in extracted RNA from PAXgene tubes has been already described *(50)*. This non-complete depletion of erythrocytes during RNA extraction might hamper to perform reliable profiles of gene expression by masking relevant genes in the white blood cells content. In order to overcome that, we decided to use an approach developed by Krjutškov K, et al. based on the blockage of Alpha and Beta-globin transcripts, two of the most highly expressed transcripts in erythrocytes *(51)*. Briefly, ZNA-modified blocking oligos (lacking 3’-OH) annealing at 3’-end of mRNA molecules after RNA denaturation prevented the binding site of oligo-T primer, and therefore the generation of cDNA from these transcripts. Twenty nanograms of total RNA were used for blocking and cDNA synthesis with SMART-seq2 method *(52)*. After PCR, DNA was inspected with Qubit dsDNA HS Assay Kit (ThermoFisher, cat.nr. Q32854) and High Sensitivity DNA Analysis Kit (Agilent Technologies, cat.nr. 5067-4626). Nextera XT was then used to prepare DNA libraries for Illumina NovaSeq 6000 S2 (2×50bp).

### OCR Measurement

Seahorse XFe96 Analyzer (Seahorse Bioscience) was used to measure oxygen consumption rate of ME-patient derived PBMCs. Frozen PBMC were recovered in RPMI 1640 medium supplemented with 1% of penicillin-streptomycin solution and 10% FBS for 4.5 hours. One hour before the assay, the growth medium was replaced with Seahorse XF RPMI medium, and 3×10^5^ PBMC per well were seeded with 40 μl assay medium in XF 96-well cell culture microplate coated with poly-D-lysine (Sigma). The plate was centrifuged at 200 x g for 1 sec with no brake, rotated 180 ° and centrifuged again for 1 sec with no brake at 300 x g. After centrifugation,140 μl of assay medium was added per well and the plate was left to stabilize in a 37°C non-CO_2_ incubator. The wells were sequentially injected with 12.64 μM oligomycin (Sigma Aldrich 75351), 20 μM FCCP (Sigma Aldrich C2920), 5 μM rotenone (Sigma Aldrich R8875) together with 5 μM antimycin A (Sigma Aldrich A8674) and the oxygen consumption rate (OCR) was measured for each well three times, every 3 min, before and after each injection. OCR was normalized to protein concentration following BCA assay conducted according to the manufacturer’s instructions.

### Mass Cytometry Preprocessing and Gating

All FCS-files were exported without any pre-processing from the CyTOF software and normalized using an in-house version of normalization software *(53)*. Debarcoding of each file was done using the MATLAB version of the single cell debarcoder *(54)*. CellGrid, a supervised learning algorithm based on t-SNE implementation, was used to gate sub-cell populations (Chen, manuscript in prep.).

### ProSeek plasma protein data preprocessing

Olink’s arbitrary unit is Normalized Protein eXpression (NPX) which is Cq-values for each protein that are recalculated to a relative log_2_ scale. The data is normalized in order to minimize intra- and inter-assay variation, and the data from the three panels were merged to one dataset. If proteins were detected in less than 20% of all samples were removed and any missing values were set to the lowest detectable value for that protein. Due to the single batch experiment no batch correction was necessary due to Olink’s built-in quality control across panels, but batch effect check was performed indicating the same conclusion.

### RNAseq data analysis

Quality control was provided by the National Genomics Infrastructure (NGI) at Science for Life Laboratory, Stockholm, Sweden. The first step after mRNA-sequencing was quantifying abundances of transcript sequences in FASTA format by generating abundance estimates for all samples using the Kallisto software *(55)*. Also, gene abundance estimates were performed by summing the transcript expression (TPM) values for the transcripts of the same gene. Since DESeq2 expects count data, from the Kallisto output the *tximport* package was used to convert these estimates into read counts. DESeq2 was performed as a basis for differential gene expression analysis based on the negative binomial distribution *(56)*. Low gene counts (<100) were filtered out and variance stabilizing transformation (VST) was performed on the count data, as well as batch correction using the *limma* package.

### Analysis of variance in Metabolomic Data (Mito stress Seahorse assay)

Mixed-effects analysis using Sidak’s multiple comparisons test over all timepoints (number of active treatments) was performed for maximal respiration and spare respiratory capacity results. Batch effect correction was performed beforehand using the *limma* package, to then perform a mixed effect model to account for paired sampling over treatment.

### Automated Cell classification

Grid is an in-house supervised algorithm based on the use of manually gated cell sub-populations as reference to train a classifier algorithm that can then classify similar cells quickly and accurately. The following populations were gated: B cells (IgD^+^ memory B cells, IgD^-^ memory B cells, naïve B cells, transitional B cells and plasmablasts), CD4 T-cells (central memory CD4T, effector memory CD4T, naïve CD4T, naïve Tregs, and memory Tregs), CD8 T-cells (activated CD8T, central memory CD8T, DP T-cells, effector memory CD8T, and naïve CD8T), eosinophils, MAIT, monocytes (classical, non-classical and proinflammatory monocytes), natural killer (NK) cells (CD56^bright^ NK and CD56^dim^ NK), neutrophils, basophils, gdT (CD161^+^ gdT and CD161^-^ gdT) and pDC. These sub-populations were identified by phenotypic markers from the parameter selection.

### Mixed-Effects Modeling

Complex mixed-effect models tend to result in singular fits; therefore, it is recommended to use a partially Bayesian method. The *blme* package was applied on this data to produce maximum a posteriori (MAP) estimates. This provided the ability to not only nest the variables, link individuals into sets of Pre- and Post-INMEST treatment, but also to account for sex, age as well as symptom scores and active treatments *(57)*.

### Multi-Omics Factor Analysis (MOFA)

The input to MOFA is a set of matrices with dimensions (plasma protein expression, cell abundance, and gene expression), the *MOFAobject* was built using *MultiAssayExperiment* object which was subsequently trained in R instead of Python through the *reticulate* package. Prior to model training as the Gaussian noise model was used, methods like DESeq2 was performed on the mRNA-seq data for normalization and variance stabilization. All data sets were processed individually to remove any features resulting in zero or low variance before fitting the model. Convergence was assessed since MOFA is trained using variational Bayes which consists of the Evidence Lower Bound (ELBO), therefore the change in ELBO (deltaELBO) was used and fit the convergence threshold which is considered to be between 1 to 10. The *MOFAobject* was trained with 10 factors and a variance threshold of 0.01%. Multiple models under different initializations were run to make sure that factors were consistently found across trials for model selection.

### Gene Set Enrichment Analysis (GSEA)

Gene set enrichment analysis (GSEA) was performed to investigate enriched gene sets in accordance to number of active treatments *(58)*. These groups of genes were derived from the mixed effect modeling and then *gseGO* was used for GSEA looking at GO terms for Biological Processes with p-values calculated from mixed effect modeling.

### Disease Tolerance Networks

Disease Tolerance Networks were built by finding key regulators for query genes using Martins et al. (*40*) from TTRUST v2. Edge and node dataframes were built from information acquired from TTRUST v2 with the addition of calculated log_2_(treated/baseline) for color gradient. Node sizes were based on number of edges, the larger the node the more edges. These were used to build the networks with *visNetwork()*.

## General

We are grateful for all patients taking part in this study, and all colleagues at Stora Sköndal ME/CFS clinic.

## Author contributions

P.J, JE.J and P.B designed the study. Patients were treated by either of two otorhinolaryngologists, of which JE.J was one and evaluated by PJ. L.R analyzed all data with support from Y.C and P.B. C.P, T.LK, J.M, J.W, C.M helped in sample collection and processing. J.Z performed immune cell metabolic experiments with support from J.R. P.B and L.R wrote the manuscript with input from all co-authors.

## Data and materials availability

FCS files from Mass Cytometry experiments available at http://flowrepository.org/id/FR-FCM-Z2F2. All correspondence and material requests should be addressed to Petter Brodin.

## Supplementary Materials

**Table S1.** List of antibodies used in mass cytometry.

| **Metal Tag** | **Antigen** | **Clone** | **Vendor** |
| --- | --- | --- | --- |
| Y89 | CD45 | HI30 | Fluidigm |
| In113 | HLA-ABC | W6/32 | BioLegend |
| In115 | CD57 | HCD57 | BioLegend |
| La139 | TCR Vα7.2 | 3C10 | BioLegend |
| Nd142 | CD19 | HIB19 | Fluidigm |
| Nd143 | CD5 | UCHT2 | BioLegend |
| Nd144 | CD16 | 3G8 | BioLegend |
| Nd145 | CD4 | RPA-T4 | BioLegend |
| Nd146 | CD8a | SK1 | BioLegend |
| Sm147 | CD11c | Bu15 | Fluidigm |
| Nd148 | CD31 | WM59 | BioLegend |
| Sm149 | CD25 | 2A3 | Fluidigm |
| Nd150 | CD64 | 10.1 | BioLegend |
| Eu151 | CD123 | 6H6 | BioLegend |
| Sm152 | $\gamma\delta$TCR | 5A6.E9 | Fischer S |
| Eu153 | CD13 | WM15 | BioLegend |
| Sm154 | CD3e | UCHT1 | Fluidigm |
| Gd155 | CD7 | CD7-6B7 | BioLegend |
| Gd156 | CD26 | BA5b | BioLegend |
| Gd157 | CD9 | SN4 C3-3A2 | eBio |
| Tb159 | CD22 | HIB22 | BioLegend |
| Gd160 | CD14 | M5E2 | BioLegend |
| Dy161 | CD161 | HP-3G10 | BioLegend |
| Dy162 | CD29 | TS2/16 | BioLegend |
| Dy163 | HLA-DR | L243 | BioLegend |
| Dy164 | CD44 | BJ18 | BioLegend |
| Ho165 | CD127 (IL-7Rα) | A019D5 | Fluidigm |
| Er166 | CD24 | ML5 | BioLegend |
| Er167 | CD27 | L128 | Fluidigm |
| Er168 | CD38 | HIT2 | BioLegend |
| Tm169 | CD45RA | HI100 | Fluidigm |
| Er170 | CD20 | 2H7 | BioLegend |
| Yb171 | CD33 | WM53 | BioLegend |
| Yb172 | IgD | IA6-2 | BioLegend |
| Yb173 | CD56 | HCD56 | BioLegend |
| Yb174 | CD99 | HCD99 | BioLegend |
| Lu175 | CD15 | W6D3 | BioLegend |
| Yb176 | CD39 | A1 | BioLegend |
| Ir191 | Cell-ID™ Intercalator-Ir (DNA) | NA | Fluidigm |
| Ir193 | Cell-ID™ Intercalator-Ir (DNA) | NA | Fluidigm |
| Bi209 | CD11b | Mac-1 | Fluidigm |

